# A heat-assisted tt*Lb*Cas12a workflow incorporating crRNA 3’-tailing enables efficient multiplex biallelic genome editing in hybrid poplar

**DOI:** 10.64898/2026.10.21.760723

**Authors:** Virginia Zahn, Alexander Fendel, Matthias Fladung, Tobias Bruegmann

## Abstract

In forest trees, high genome-editing efficiency in primary transformants is particularly important for functional genomics and trait improvement, as long generation times severely limit the use of subsequent generations to recover fully edited genotypes. This challenge is further compounded in hybrid poplar, where paralogous gene copies and allelic variation make efficient multiplex editing particularly difficult. We optimized *Lb*Cas12a-mediated editing in *Populus* × *canescens* (*Populus tremula* × *Populus alba*), clone INRA 717-1B4, by targeting four paralogous *PtaABA8ox4* loci using temperature-tolerant tt*Lb*Cas12a, a two-stage heat treatment, and crRNA 3′-tailing. The heat-treatment regime preserved transgenic recovery, while *ttLbCas12a*-based constructs and heat treatment substantially increased biallelic editing. All seven lines in the highest-performing group, combining tt*Lb*Cas12a, 3′-tailed crRNAs, and heat treatment, were biallelically edited at all four targets. crRNA 3′-tailing altered predicted secondary structures and showed a consistent numerical tendency toward improved editing, but its independent effect was not statistically resolved. Mutations were dominated by small deletions, and biallelic out-of-frame recovery increased in *ttLbCas12a*-based contexts without scaling proportionally with biallelic editing. Thus, heat-assisted tt*Lb*Cas12a-mediated approaches enable efficient multiplex editing in hybrid poplar, whereas genotype quality remains dependent on repair outcome and somatic uniformity.

## 1 Introduction

Trees of the genus *Populus* (poplars, aspens, and cottonwoods) are among the world’s most important fast-growing trees and hold global significance for wood production, biomass yield, carbon sequestration, and climate change mitigation (Böhlenius et al. 2023; Enescu et al. 2026). Owing to their rapid growth, the availability of genomic data for several clones, efficient genetic transformation, and *in vitro* regeneration, poplars are frequently used as model organisms for biotechnology research among trees (Thakur et al. 2021; Bruegmann et al. 2024). Genetic modification—commonly performed via *Agrobacterium tumefaciens*—is a well-established tool for functional studies and trait improvement in *Populus* (Leple et al. 1992; Wang et al. 2025). Given the central role of genetic modification in elucidating gene function and improving breeding- and production-relevant traits in poplar, CRISPR/Cas systems have become indispensable tools (Ahuja 2021; Cao et al. 2022). CRISPR/Cas-mediated genome editing—particularly by Cas9 or Cas12a—enables targeted DNA double-strand breaks (DSBs), facilitating both gene knockouts (KOs) and more sophisticated modifications (Jinek et al. 2012; Mao et al. 2013; Zetsche et al. 2015; Wang et al. 2017).

Despite this comparatively high experimental tractability, genome editing in *Populus* spp. remains challenging due to the genomic complexity of this genus. This is particularly relevant for interspecific hybrid clones such as the widely used INRA 717-1B4 (*Populus tremula* × *Populus alba*), which is highly amenable to *Agrobacterium*-mediated transformation and *in vitro* regeneration but combines extensive allelic sequence variation between its parental genomes with numerous paralogous gene copies retained from ancient genome duplications (Tuskan et al. 2006; Mader et al. 2016; Bruegmann et al. 2019; Li et al. 2023b). When simultaneous disruption of several functionally relevant gene-family members is required, paralogy increases the number of loci that must be edited and may therefore require multiple highly efficient, target-determining CRISPR-RNAs (crRNAs). Sequence divergence between parental alleles can impair simultaneous targeting of both alleles through spacer or protospacer-adjacent motif (PAM) mismatches (Kim et al. 2017), further increasing the number of required crRNAs or restricting guide design to target sites conserved across both alleles. In addition, allelic sequence variation within or around cleavage sites may contribute to allele-dependent differences in editing efficiency and repair outcomes, potentially preventing complete loss of gene function if one allele remains unedited or retains an in-frame mutation (Kim et al. 2017; Bernabé-Orts et al. 2019; Lageix et al. 2024). In parallel, high primary editing efficiencies are required to recover complete multiplex editing directly in primary transformants. Incomplete editing of individual loci or alleles would otherwise require additional rounds of transformation and editing or the fixation and combination of mutations through subsequent sexual generations. Re-transformation would substantially prolong the experimental workflow, whereas genetic segregation is particularly impractical in poplar because natural flowering generally begins only after approximately 7–10 years of juvenile growth (Hsu et al. 2006; Cao et al. 2022; Bruegmann et al. 2024).

While genome editing utilizing Cas9 is well established in *Populus*, Cas12a has only been applied in a few initial studies (An et al. 2020; Li et al. 2023a; Zhang et al. 2023; Das et al. 2025). In contrast to Cas9, Cas12a offers distinct advantages: it requires a TTTV PAM, thereby unlocking AT-rich genomic regions. It generates staggered DSBs distal to the PAM, which are frequently associated with deletion-dominated repair outcomes and can thereby increase the likelihood of disruptive mutations, and its compact crRNA architecture facilitates multiplex targeting (Zetsche et al. 2015; Wang et al. 2017; Bernabé-Orts et al. 2019). However, achieving high editing efficiencies by Cas12a in poplars remains challenging (An et al. 2020; Das et al. 2025). Several factors can constrain Cas12a-mediated genome editing in plants, including (i) low Cas nuclease activity at standard tissue-culture temperatures; (ii) chimerism resulting from delayed editing events; and (iii) suboptimal crRNA design (Kim et al. 2017; Malzahn et al. 2019; Wang et al. 2021; Li et al. 2023a; Ren et al. 2023). Moreover, prolonged or asynchronous editing may increase somatic genotype heterogeneity (chimerism) and thereby complicate the regeneration of genetically uniform edited lines.

To overcome these constraints, Cas12a-mediated editing can be optimized at several levels. Temperature sensitivity and delayed editing can be addressed by enzyme engineering, including the temperature-tolerant *Lachnospiraceae bacterium* Cas12a variant tt*Lb*Cas12a (D156R) with enhanced activity at typical plant cultivation temperatures, and by regeneration protocols incorporating early heat treatments to promote nuclease activity shortly after transformation (LeBlanc et al. 2018; Merker et al. 2020; Schindele and Puchta 2020; Zhang et al. 2023). In addition to editing activity itself, the selection regime can influence which transformed events are recovered during regeneration, as stronger selection preferentially retains cells with sufficient selection marker expression to survive and regenerate (Zhou et al. 2013). Finally, editing performance may be improved by optimizing crRNA structure. Multiplexing with several crRNAs increases the chance of recovering edited alleles but can be constrained by target-site availability, positional effects, crRNA folding, and vector construction complexity (Lowder et al. 2015; Wang et al. 2017; Creutzburg et al. 2020; Ren et al. 2023). Alternatively, adding nucleotides to the 3--end of crRNAs that are not complementary to the target sequence has been proposed to enhance editing by reducing unfavorable intramolecular spacer interactions and promoting formation of the functional crRNA pseudoknot (Creutzburg et al. 2020).

Building upon these strategies, this study aims to systematically optimize *Lb*Cas12a-mediated genome editing in the poplar hybrid clone INRA 717-1B4. To demonstrate the feasibility of our approach, we targeted the *ABSCISIC ACID 8′-HYDROXYLASE 4* (*PtaABA8ox4*) gene family as a proof-of-concept system, aiming to simultaneously edit all four paralogs as a stringent test of multiplex genome editing. The four *PtaABA8ox4* paralogs are predicted to encode ABA 8′-hydroxylases of the CYP707A family, whose members catalyze a key step in oxidative ABA catabolism (Kushiro et al. 2004). Thus, PtaABA8ox4 provided both a biologically relevant stress-related target for further research, without aiming to resolve its physiological consequences in the present study, and a stringent technical test case for multiplex editing in a heterozygous woody species. To this end, we implemented three optimization strategies: (i) comparison of wild type (WT) *Lb*Cas12a with the temperature-tolerant tt*Lb*Cas12a variant, (ii) application of a heat treatment protocol, and (iii) evaluation of crRNA 3′-tailing. We first evaluated transgenic recovery under the two selection regimes and heat-treatment conditions, then systematically evaluated the effects of CRISPR/Cas construct, heat treatment, and target identity on general editing and biallelic editing performance, followed by focused analyses within the shared tt*Lb*Cas12a background to determine whether crRNA 3′-tailing altered target-specific editing performance. Finally, we characterized allele specificity, chimerism, deletion patterns, and predicted reading-frame consequences.

## 2 Materials and Methods

### 2.1 Plant Material

Hybrid poplar INRA 717-1B4 (*P. tremula* × *P. alba*; Leple et al. (1992)) was maintained *in vitro* on shoot elongation medium (SEM), based on Woody Plant Medium (WPM; Lloyd and McCown (1980)) with 2% sucrose and 0.65% agar, pH 5.8, at 22 °C under a 24-h photoperiod (approx. 30 µmol m^−2^ s^−1^).

### 2.2 Identification of *PtaABA8ox4* Homologs

The rice *Os*ABA8ox4 protein (LOC_Os09g28390; Zhang et al. (2020)) was queried by BLASTp against *P. tremula* × *P. alba* HAP1/HAP2 v5.1 at Phytozome (Goodstein et al. 2012; Zhou et al. 2023). Candidates with ≥ 67% amino-acid identity were retained. Four loci on chromosome (chr) 2, 4, 9, and 14 comprised the HAP1/HAP2 pairs PtXaTreH.02G107700 / PtXaAlbH.02G108600, PtXaTreH.04G110400 / PtXaAlbH.04G109700, PtXaTreH.09G084200 / PtXaAlbH.09G086000, and PtXaTreH.14G020700 / PtXaAlbH.14G019600, respectively, and were designated *PtaABA8ox4* targets. Based on sequence homology and chromosomal distribution, these loci were assigned to two paralogous pairs: chr 2 / chr 14 and chr 4 / chr 9.

### 2.3 Design and Optimization of Cas12a crRNAs

Target-site and crRNA design followed Zahn et al. (2026). crRNA1 targeted both alleles of chr 2/14 and crRNA2 those of chr 4/9. Optimized variants carried four additional 3′ nucleotides selected to improve predicted crRNA folding while avoiding extensive, but not all, tail–target complementarity (Creutzburg et al. 2020). Spacer GC content was 45–50% (Table S1). Centroid and minimum-free-energy (MFE) structures were predicted with RNAfold (Lorenz et al. 2011) at 22 and 37 °C using Andronescu et al. (2007) parameters and including an additional 3′ U representing a potential transcript residue derived from the downstream 8×T Pol III terminator. Because the single lowest-energy MFE structure was not representative of the predicted ensemble for most crRNAs, centroid structures were used as ensemble-representative folds for subsequent structural comparisons. Cas-OFFinder screened the INRA 717-1B4 genome for ‘TTTN’ PAM sites with ≤ 3 mismatches and no bulges (Bae et al. 2014).

### 2.4 Construction of CRISPR/Cas12a Vectors

Three constructs were synthesized by DNA Cloning Service (Hamburg, Germany). The vector ‘ABA-Cas’ contained the WT variant of *LbCas12a*, crRNA1, crRNA2, and the hygromycin-resistance gene *HYGROMYCIN PHOSPHOTRANSFERASE II* (*HPTII*). ‘ABA-ttCas’ contained *ttLbCas12a*, crRNA1, crRNA2, and the kanamycin-resistance gene *NEOMYCIN PHOSPHOTRANSFERASE II* (*NPTII*). ‘ABA-ttCas.opt’ contained *ttLbCas12a*, crRNA1.opt, crRNA2.opt, and *NPTII*. *HPTII* and *NPTII* expression was driven by the *A. tumefaciens NOPALINE SYNTHASE* (*NOS*) promoter, *LbCas12a*/*ttLbCas12a* expression by the *Arabidopsis thaliana UBIQUITIN10* promoter, and each crRNA by a separate *A. thaliana U6-26* (*AtU6*-*26*) promoter (Horstmann et al. 2004; Li et al. 2007; Zhou et al. 2013; Wolabu et al. 2020). Fully annotated vector sequences are provided in Data S1. Constructs were introduced into *A. tumefaciens* strain GV3101-pMP90 by heat shock (Höfgen and Willmitzer 1988).

### 2.5 *Agrobacterium*-mediated Transformation

Poplar leaf and stem transformation followed the protocol by Bruegmann et al. (2019) with modifications. Explants were co-cultivated with *A. tumefaciens* for 3 d on SEM at 22 °C (–HT) or 28 °C (+HT regime), then transferred to 150 mL liquid shoot-induction medium (SIM: WPM, 2% sucrose, 100 mg/L Pluronic F-68, 2.2 µg/L TDZ, pH 5.8) containing 500 µg/mL cefotaxime and either 20 µg/mL hygromycin (ABA-Cas) or 50 µg/mL kanamycin (ABA-ttCas/ABA-ttCas.opt). Cultures were shaken at 100 rpm for four weeks in darkness, at 22 °C or under the +HT regime. Cultures were then de-etiolated for two weeks in dim light, transferred to solid SIM (0.65% agar) for shoot-cluster formation, and individual shoots (one per cluster) were moved to ½ SIM containing 1.1 µg/L TDZ for elongation. All the medium ingredients were purchased from Duchefa (Haarlem, The Netherlands).

### 2.6 Heat Treatment Strategy

The +HT regime combined 28 °C co-cultivation, intended to support T-DNA transfer (Movahedi et al. 2014), with cyclic 37 °C treatment during regeneration to enhance (tt)*Lb*Cas12a activity (Schindele and Puchta 2020); adapted from LeBlanc et al. (2018). A pilot study comparing treatment starts at 3 d and 1, 2, or 3 weeks identified one week after co-cultivation as the earliest compatible with shoot vitality. Four 30h-cycles at 37 °C were then applied, separated by 42 h at 22 °C. All heat-treatment cycles and intervening recovery periods were conducted in darkness. For ABA-Cas and ABA-ttCas, transformations with and without heat treatment were performed in parallel to evaluate potential effects of the temperature regime on regeneration and to facilitate comparison of the *HPTII*- and *NPTII*-based selection systems.

### 2.7 Verification of Transgenic Lines and Target-site Amplification

Genomic DNA was isolated according to Bruegmann et al. (2022). Lines PCR-positive for both the Cas nuclease and *HPTII*/*NPTII* were classified as transgenic. Transgenic recovery was calculated for each transformation as the proportion of regenerated lines confirmed as transgenic by PCR. To limit strongly unequal sample sizes among construct × heat-treatment groups, PCR-confirmed lines were randomly subsampled for four-target genotyping independently of subsequent genotype. Gene-specific PCR amplified each *PtaABA8ox4* target. Ambiguous heterozygous chromatograms were resolved by allele-specific PCR using diagnostic SNPs (A1 = *P. tremula*; A2 = *P. alba*). Primers are listed in Table S2.

### 2.8 Sequence Analysis and Genotype Classification

PCR products were Sanger sequenced (StarSEQ, Mainz, Germany) and aligned to reference sequences in SnapGene. Alleles were assigned as A1/A2 using diagnostic SNPs. Overlapping peaks in allele-specific reads at or downstream of the expected cleavage site indicated somatic genotypic heterogeneity (Figure S1). Because genotyping could not distinguish heterogeneity arising from multicellular origin (chimerism) from that caused by ongoing editing within a common lineage (mosaicism), “chimerism” was used here as an umbrella term for both. A target was scored edited only when at least one allele was fully edited with no remaining WT sequences. Genotypes were classified as WT, monoallelic, biallelic heterozygous, or biallelic homozygous, with chimerism recorded separately.

Manual chromatogram evaluation provided primary genotype calls. A custom R workflow additionally derived mutation classes and deletion coordinates for positional, recurrent-product, and recognition-site analyses (Data S2). Reads were aligned in both orientations to target-specific references, with mutation calling restricted to a 21 bp window centered on the cleavage site. The workflow was developed on synthetic data and evaluated against a separate synthetic test set not used during workflow development (hold-out set), as well as manually annotated experimental reads. Breakpoints were also blindly re-annotated allowing sequence-equivalent coordinates. Full algorithms and validation are described in Method S1.

### 2.9 Quantification of Editing Performance and Mutation Patterns

Performance definitions and observation levels are summarized in Table 1. Chimerism was quantified separately among biallelic edits. Full biallelic editing required both alleles edited at all four loci. Mutation class (deletion, insertion, or mixed InDel) was assigned per allele, with deletion analyses restricted to pure deletions ≥ 1 bp. Recognition-site retention was categorized by PAM disruption and contiguous PAM-proximal spacer sequence (≤ 16, 17–18, or ≥ 19 nt), reflecting reported Cas12a complementarity thresholds (Naqvi et al. 2022; Ren et al. 2023). Parental-allele analyses used monoallelic observations, with A1/A2 retained descriptively for deletion analyses. Indels divisible by three were classified in-frame. All others were out-of-frame (OOF), and biallelic OOF required OOF edits in both alleles. Positional deletion profiles were calculated as the proportion of pure-deletion alleles deleting each reference position. Recurrent deletion products were grouped by deletion coordinates within each target.

**Table 1:**
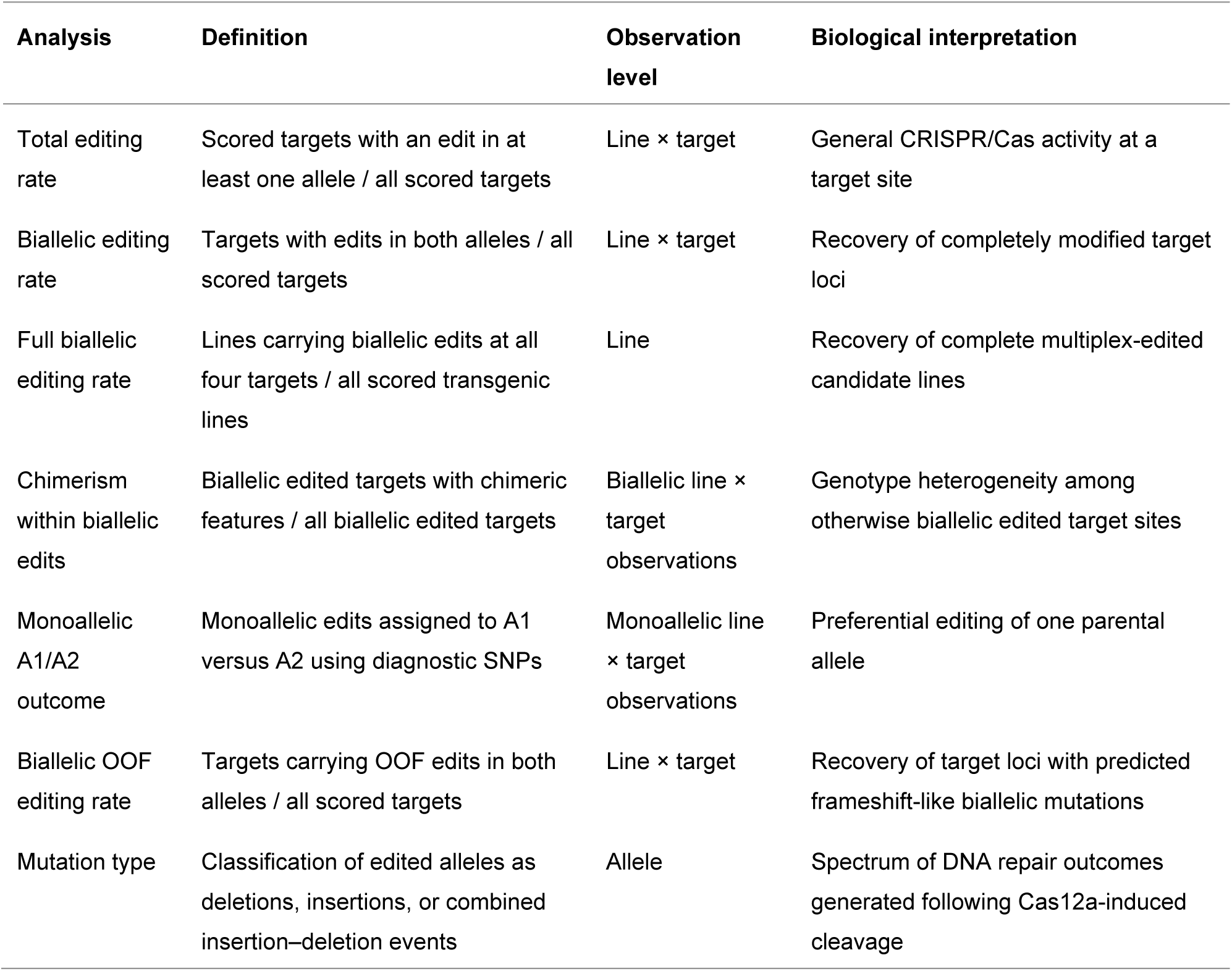
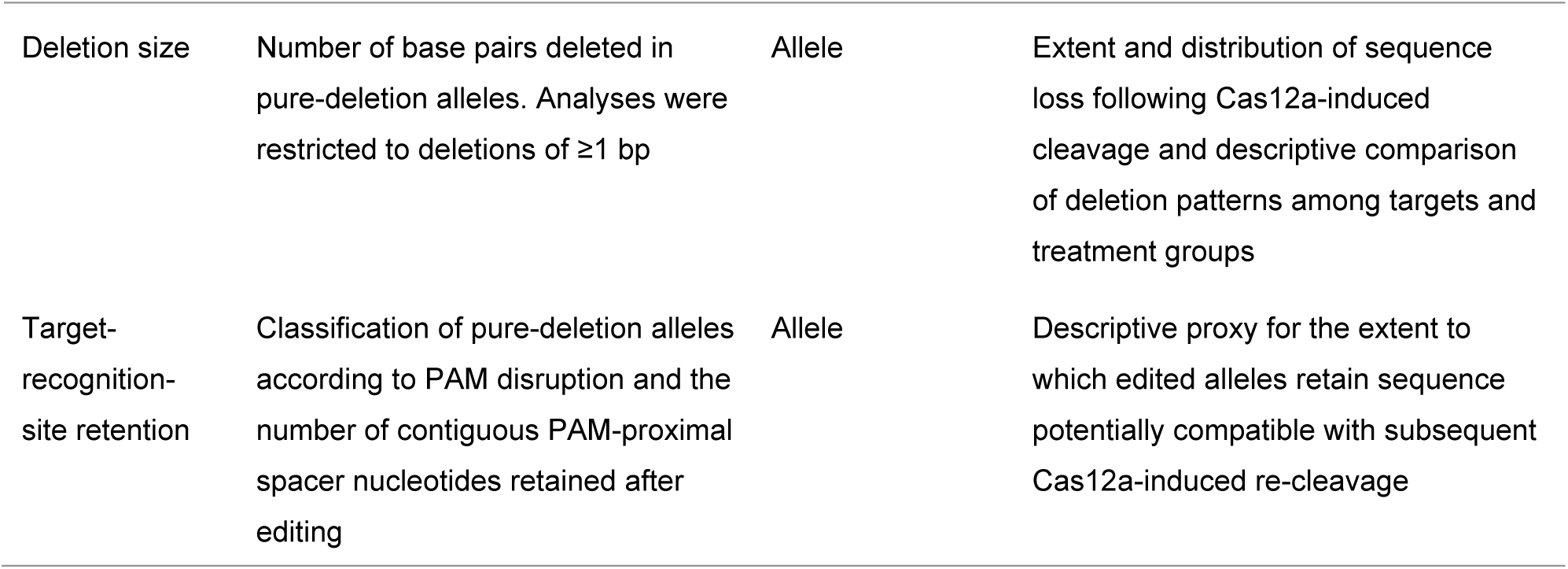
Definitions and observation levels of genome-editing analyses used in this study.

### 2.10 Statistical Analysis and Data Visualization

Raw genotype and PCR data are in Data S3. Analysis scripts are available at Zenodo (Zahn 2026) graphical data in Data S4, and full statistical outputs in Data S5. Analyses used R 4.6.0 (R Core Team 2026) with glmmTMB, emmeans, DHARMa, dplyr, tidyr, ggplot2, and ggpattern. Target-level binary outcomes (total editing, biallelic editing, chimerism within biallelic edits, monoallelic A1/A2 outcome, and biallelic OOF editing) were analyzed by binomial-logit generalized linear mixed models (GLMMs). Full biallelic editing and transgenic recovery were analyzed at line level. Full-dataset target models included construct, heat, and target as additive fixed effects. Construct was used as a composite factor because nuclease variant and crRNA optimization were not fully crossed. Thus, each fixed effect represented its average association with the response after adjustment for the other fixed effects. No fixed-effect interaction terms were included. Random intercepts were included for transformation, transformation × target, and line. Here, transformation accounted for shared experimental origin, transformation × target for transformation-specific variation among targets, and line for repeated target-level observations within a regenerated line. Transgenic recovery was analyzed separately using selection marker and heat as fixed effects and transformation as a random intercept. The focused ABA-ttCas/ABA-ttCas.opt analysis tested the crRNA optimization × target interaction—whether the optimization effect differed among loci— with heat retained as an additive adjustment. Prespecified fixed effects were assessed by likelihood-ratio tests (LRTs) and 999-simulation parametric-bootstrap LRTs (PB-LRTs). Estimated marginal means (EMMs) used Tukey adjustment for full-dataset multi-level comparisons, while focused optimized-versus-non-optimized contrasts were Holm-adjusted and within-condition target comparisons Tukey-adjusted. WT-chimeric calls were descriptive. DHARMa diagnostics showed no major deviations (Figures S2, S3; Table S3). Detailed model specifications and robustness analyses are in Method S2, Data S5.

## 3 Results

### 3.1 *ABA8ox4* target identification and predicted effects of 3’-tailing on crRNA folding

BLASTp against the *P. tremula* × *P. alba* genome identified eight haplotype-specific protein models representing four ABA8ox4 loci with 67–71% amino-acid identity to rice *Os*ABA8ox4 (Table S4). CrRNA1/crRNA1.opt targeted the loci on chr 2 and chr 14 while crRNA2/crRNA2.opt targeted loci on chr 4 and chr 9 (Figure 1A). RNAfold centroid structures indicated that 3′-tailing reduced spacer self-pairing in crRNA1.opt while retaining the pseudoknot-associated pattern, whereas crRNA2.opt showed increased pairing probability in the pseudoknot-forming region (Figure 1B). Tail–target complementarity was limited to crRNA2.opt, which showed allele-specific complementarity at chr 9 (Table S1). Potential allele bias was therefore examined in section 3.3.2. Cas-OFFinder detected no chromosome-assigned off-targets. One low-priority crRNA2 candidate with a TTTT PAM and three mismatches occurred on an unanchored scaffold. Predictions at 37 °C retained the same tailing-associated structural differences observed at 22 °C (Figure S4).

**Figure 1:**
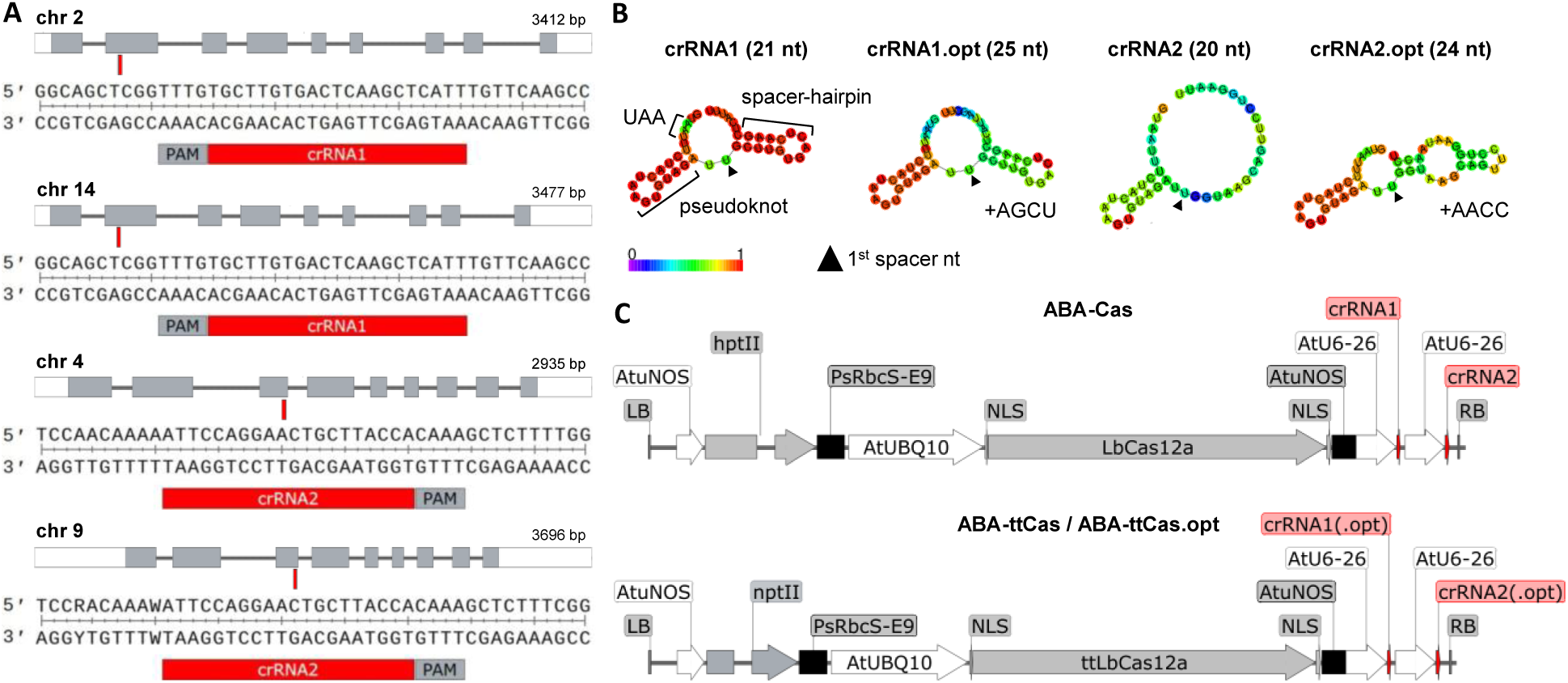
CRISPR targets, crRNA structures, and vector design. (A) Four paralogous *PtaABA8ox4* genes. Exons are grey, UTRs white, and target sites red. Insets show crRNA1/crRNA2 target sequences with PAMs in grey. (B) RNAfold centroid structures at 22 °C. +AGCU and +AACC denote 3′ tails. Colour indicates base-pair probability. Triangles mark spacer nt 1. (C) Besides the selectable-marker (*hptII*/*nptII*), the T-DNA of ABA-Cas carried *LbCas12a* and two unmodified crRNAs. ABA-ttCas and ABA-ttCas.opt carried *ttLbCas12a* with two unmodified or optimized crRNAs, respectively.

### 3.2 The heat-treatment regime preserves transgenic recovery

To identify a heat-treatment window compatible with regeneration, a pilot experiment compared treatment initiation after 3 d and 1, 2, or 3 weeks. One week after co-cultivation was the earliest 37 °C start compatible with healthy shoots, whereas earlier treatment reduced regeneration and later treatment increased stress symptoms (Figure S5). Subsequent transformation experiments comprised all three constructs (Figure 1C) under both –HT and +HT conditions (Table 2). +HT combined 28 °C co-cultivation with cyclic 37 °C treatment during regeneration (Figure 2A). Six weeks after transformation, heat treatment did not visibly impair shoot elongation. Hygromycin-selected regenerants developed more slowly but showed higher PCR-confirmed transgenic recovery than kanamycin-selected regenerants (Figure 2B, C). An additive binomial GLMM supported an effect of selection-marker system on transgenic recovery (LR = 7.99, df = 1, *p* = 0.005, PB *p* = 0.021) but no effect of the heat treatment (LR = 0.06, df = 1, *p* = 0.814, PB *p* = 0.867; Figure 2D). Despite heterogeneity among independent transformations, the selection marker-associated direction was retained across all leave-one-transformation-out refits, whereas the heat estimate varied in direction.

**Figure 2:**
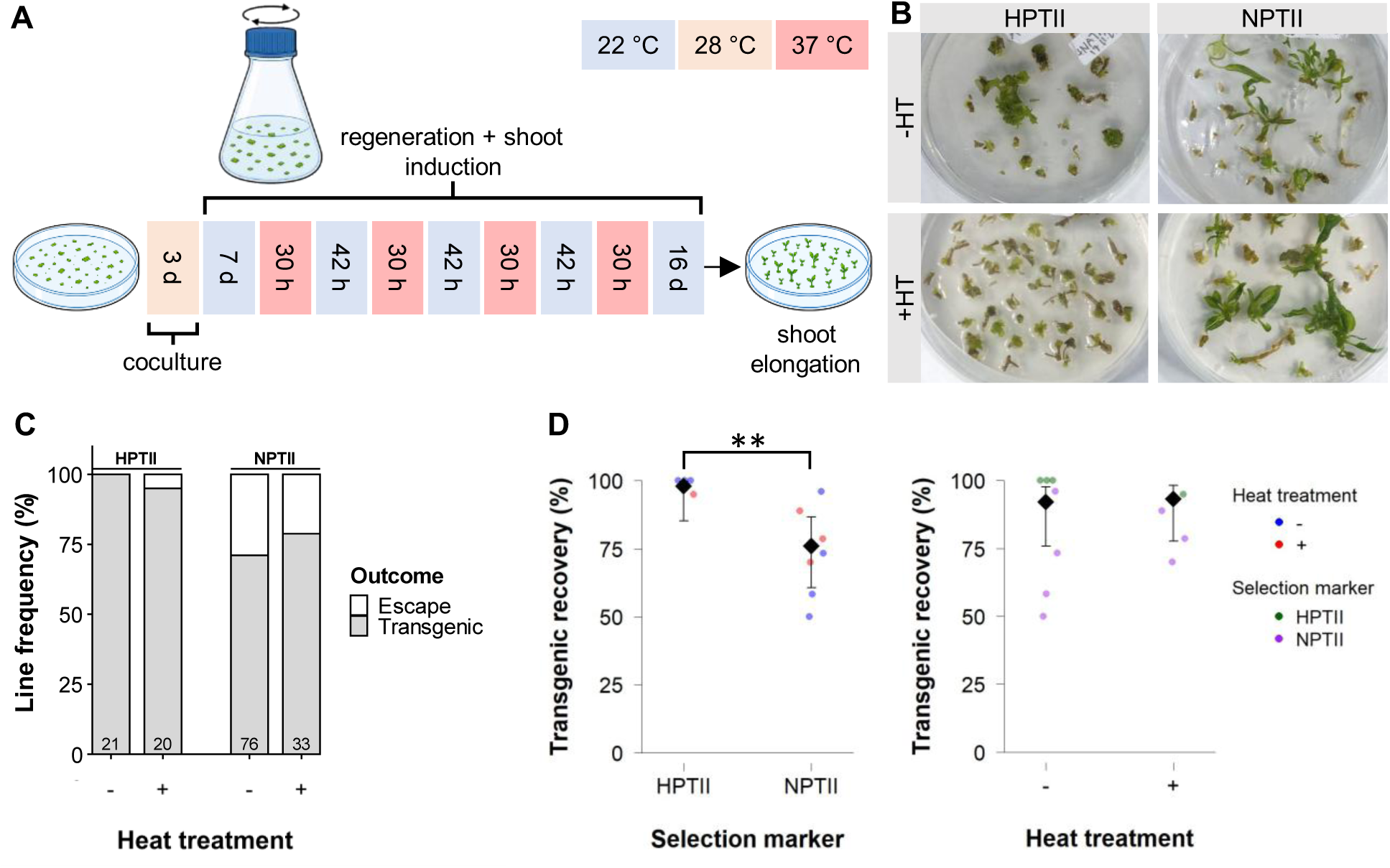
Heat-treatment regime and transgenic recovery. (A) +HT workflow: 3 d co-cultivation at 28 °C, 7 d at 22 °C, then four 30-h cycles at 37 °C separated by 42 h at 22 °C. –HT controls remained at 22 °C. All steps were performed in darkness except the initial 2 d of the 16-d regeneration phase, which were under dim light before transfer to shoot elongation medium. (B) Representative regenerants six weeks after transformation under hygromycin (HPTII) or kanamycin (NPTII) selection, either +HT or -HT. (C) PCR-confirmed transgenic and escape frequencies. Escape lines failed at least one of the two T-DNA PCR criteria. Numbers in the bars indicate total PCR-scored lines. (D) GLMM estimated marginal means (diamonds, 95% confidence intervals) for selection marker and heat treatment. Points show transformation-level observations. Asterisks indicate asymptotic LRT significance (**, *p* < 0.01).

**Table 2:**
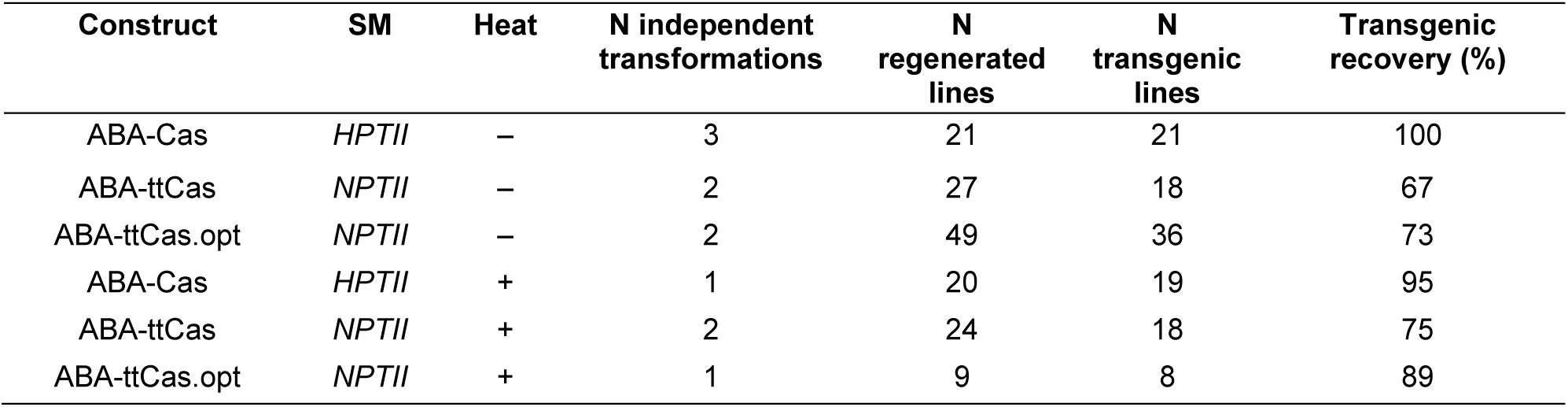
Transformation experiments by CRISPR/Cas construct, selection marker (SM), and heat treatment condition. Transgenic recovery is the proportion of regenerated lines PCR-positive for both T-DNA markers.

### 3.3 Construct design and heat treatment are associated with biallelic editing efficiency

We genotyped all four targets in 99 PCR-confirmed transgenic lines from 11 independent transformations (Figure 3A). Pooled across heat treatments and targets, ABA-Cas produced 59.2% WT and 18.4% monoallelic genotypes, whereas genotypes derived from both *ttLbCas12a*-based constructs were predominantly biallelic (60.2% and 74.1% for ABA-ttCas and ABA-ttCas.opt, respectively). Descriptively, ABA-ttCas.opt showed higher biallelic editing proportions than ABA-ttCas in all target × heat combinations except the target on chr 2 without heat treatment. Heat treatment increased the observed proportion of biallelic genotypes across all targets and constructs (Figure 3B). Total editing rate, biallelic editing rate, and full biallelic editing rate ranged from 27.5–100%, 8.8–100%, and 5.0– 100%, respectively (Table 3). Despite variation in absolute rates, experimental effects were broadly consistent across targets (Table S5). All seven heat-treated ABA-ttCas.opt lines were biallelic at all four targets (Figure 3C). Heat treatment increased all three editing measures (editing: LR = 8.33, df = 1, *p* = 0.004, PB *p* = 0.029; biallelic: LR = 10.06, df = 1, *p* = 0.002, PB *p* = 0.006; full biallelic: LR = 8.60, df = 1, *p* = 0.003, PB *p* = 0.007; Figure 3D). Construct effects were supported for biallelic editing (LR = 25.25, df = 2, *p* < 0.001, PB *p* = 0.010) and full-biallelic editing (LR = 17.01, df = 2, *p* < 0.001, PB *p* = 0.002). For total editing rate, the asymptotic LRT supported a construct effect (LR = 14.05, df = 2, *p* < 0.001), whereas the PB-LRT did not (PB *p* = 0.086). Pairwise comparisons supported higher biallelic and full-biallelic performance of both *ttLbCas12a*-based contexts than ABA-Cas, with no distinction between ABA-ttCas and ABA-ttCas.opt (Figure 3E). Target effects were not supported for editing (LR = 2.38, df = 3, *p* = 0.497, PB *p* = 0.587) or biallelic editing (LR = 2.66, df = 3, *p* = 0.447, PB *p* = 0.478; Figure S6).

**Figure 3:**
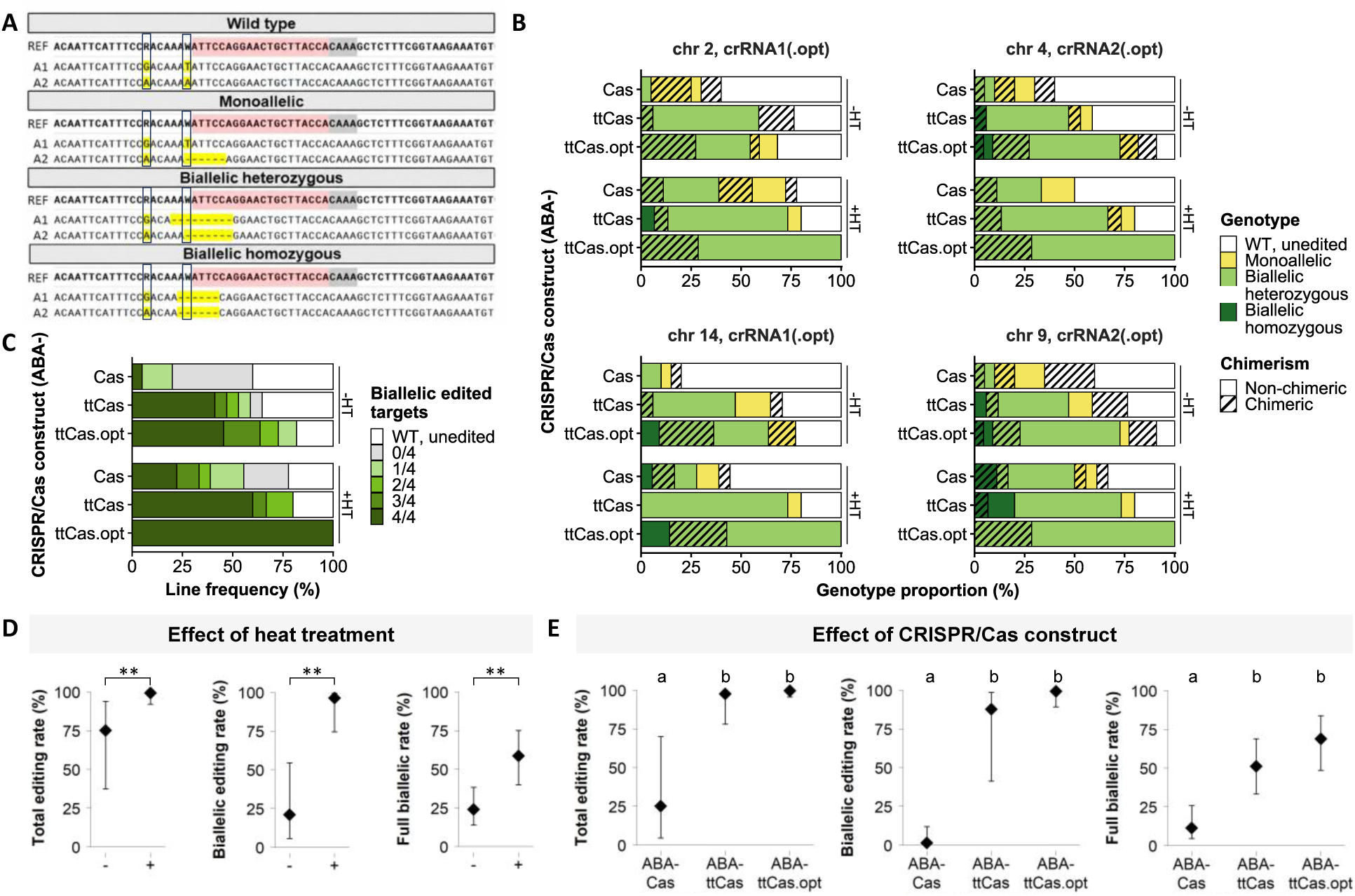
Genotypes and editing efficiency across constructs, heat treatments, and targets (99 lines, 11 transformations). (A) Representative WT, monoallelic, biallelic heterozygous, and biallelic homozygous sequences compared with the reference sequence (REF). Boxed positions mark allele-specific SNPs distinguishing the *P. tremula*-derived A1 and *P. alba*-derived A2 alleles. Yellow shading denotes CRISPR/Cas-induced changes and dashes deletions. (B) Genotype distributions by target and construct **×** heat treatment (+/-HT). Targets are arranged as crRNA1/crRNA1.opt and crRNA2/crRNA2.opt pairs to facilitate comparison of corresponding guide variants. (C) Proportions of lines carrying biallelic edits at zero to four targets. WT denotes lines without detectable edits, whereas 0/4 denotes edited lines lacking a biallelic target. (D,E) GLMM estimated marginal means (diamonds, 95% confidence intervals) for heat treatment and construct effects on editing, biallelic editing, and full-biallelic editing. Asterisks indicate asymptotic LRT significance (**, *p* < 0.01). Letters show Tukey-adjusted construct differences.

**Table 3:**
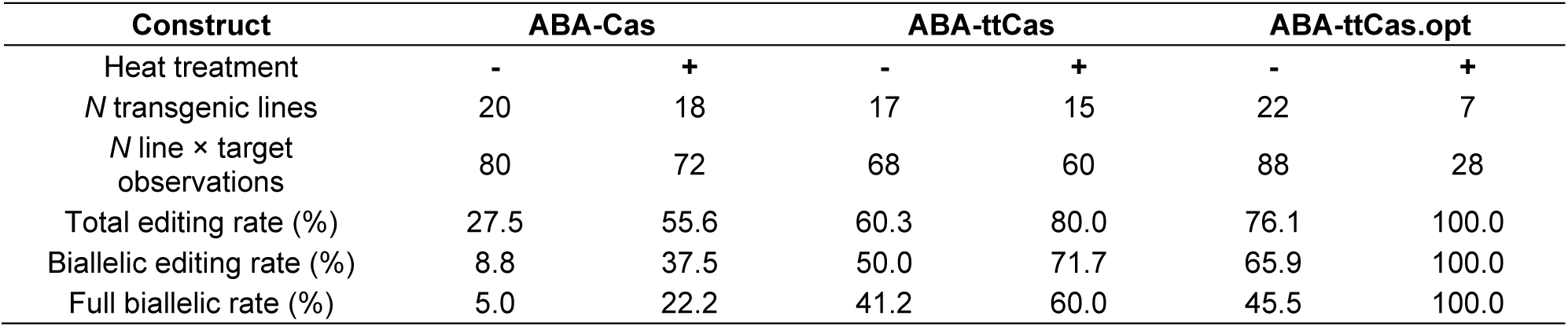
Editing outcomes pooled across targets by construct × heat treatment. WT-chimeric observations retaining WT sequence were not counted as edited.

#### 3.3.1 Increased editing is not consistently accompanied by increased chimeric biallelic outcomes

WT-chimeric calls, defined as mixed target-site signals retaining detectable WT sequence and therefore not counted as edited, occurred in 24 out of 396 observations (6.1%) and were summarized descriptively. No WT-chimeric calls occurred under heat treatment in either *ttLbCas12a* construct. Among biallelic edits, 48/197 observations (24.4%) were chimeric. Observed chimerism was lowest overall for ABA-Cas and ranged from 9.3% to 36.2% across construct × heat combinations, while target-specific frequencies were similar (22.6–26.5%; Figure S7). Construct inference was calibration-dependent (LR = 1.83, df = 2, *p* = 0.401, PB *p* = 0.019), whereas neither heat treatment (LR = 0.41, df = 1, *p* = 0.524, PB *p* = 0.092) nor target identity (LR = 2.79, df = 3, *p* = 0.424, PB *p* = 0.634) was supported.

#### 3.3.2 No systematic allele-specific editing bias was detected

Among 49 monoallelic observations, 23 involved the *P. tremula*-derived A1 allele (46.9%) and 26 the *P. alba*-derived A2 allele (53.1%). Minor target-specific trends, including at chr 9 where crRNA2.opt had partial allele-specific tail complementarity, were inconsistent (Figure S8a, b). The GLMM supported no difference in allele preference among constructs (LR = 0.44, df = 2, *p* = 0.802, PB *p* = 0.796), heat-treatment conditions (LR = 1.11, df = 1, *p* = 0.292, PB *p* = 0.316), or targets (LR = 3.79, df = 3, *p* = 0.285, PB *p* = 0.346), and no pairwise differences were detected (Figure S8c).

#### 3.3.3 Biallelic out-of-frame recovery increases in *ttLbCas12a*-based construct contexts but does not scale proportionally with biallelic editing

Biallelic OOF genotypes were generally more frequent in *ttLbCas12a*-based contexts than in ABA-Cas (Figure 4A), but the relative values for ABA-ttCas and ABA-ttCas.opt varied among targets and heat conditions, and heat-associated changes likewise differed across construct–target combinations. Thus, OOF recovery did not mirror the pattern of overall biallelic editing. Construct affected biallelic OOF editing (LR = 9.56, df = 2, *p* = 0.008, PB *p* = 0.035), with model estimates of 7.6% for ABA-Cas and 24.5–24.7% for the *ttLbCas12a* contexts. Pairwise comparisons distinguished both *ttLbCas12a*-based constructs from ABA-Cas but not from each other (Figure 4B). Target identity was not supported (LR = 2.41, df = 3, *p* = 0.492, PB *p* = 0.568), and the numerical heat-associated increase from 12.4% to 23.1% was not statistically supported (LR = 3.68, df = 1, *p* = 0.055, PB *p* = 0.105). At line level, ABA-Cas without heat treatment produced biallelic OOF edits at no more than one target, whereas the other conditions yielded lines with biallelic OOF edits at two or three targets. No line carried biallelic OOF edits at all four loci (Figure 4C), and OOF outcomes occurred at every target without obvious locus-specific depletion (Figure S9).

**Figure 4:**
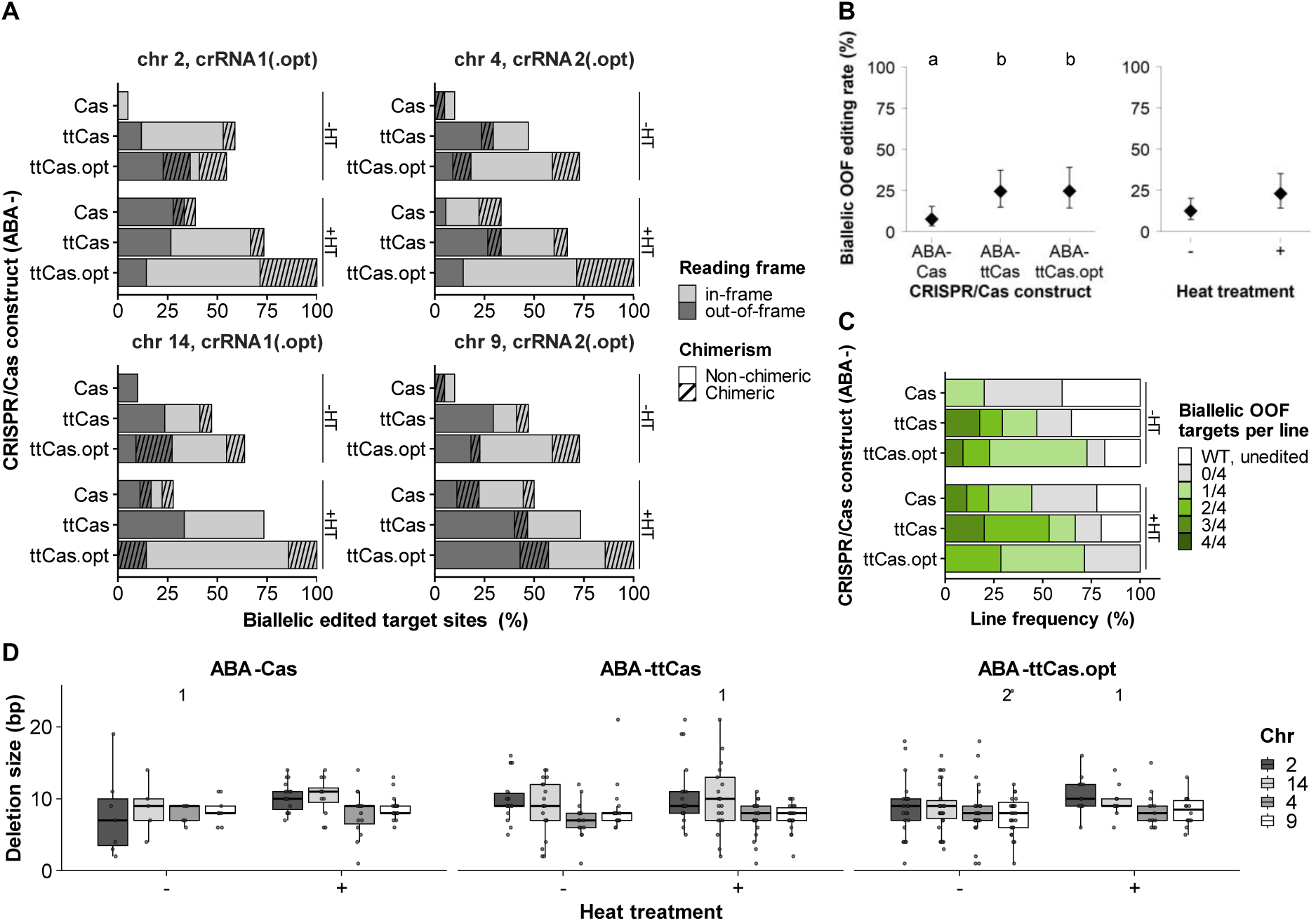
Biallelic out-of-frame (OOF) editing and deletion size. (A) Biallelic in-frame/OOF frequencies by target and treatment. Hatching denotes chimerism. (B) GLMM estimated marginal means (diamonds, 95% confidence intervals) for construct and heat effects on biallelic OOF editing. Letters show Tukey-adjusted construct differences. (C) Proportions of lines carrying biallelic OOF edits at zero to four target loci. (D) Deletion-size distributions by construct × heat and target. Boxes show medians and interquartile ranges (IQRs). The numbers above the boxes indicate the number of deletions >25 bp.

#### 3.3.4 Focused tt*Lb*Cas12a analyses reveal no consistent target-dependent effect of crRNA 3’-tailing

To test whether the effect of crRNA optimization differed among loci, we analyzed 61 lines from seven transformations within the shared *ttLbCas12a* background. The crRNA optimization × target interaction was not supported for editing in general (LR = 3.64, df = 3, *p* = 0.303, PB *p* = 0.531), biallelic editing (LR = 7.51, df = 3, *p* = 0.057, PB *p* = 0.129), chimerism (LR = 5.16, df = 3, *p* = 0.160, PB *p* = 0.302), or biallelic OOF editing (LR = 2.92, df = 3, *p* = 0.404, PB *p* = 0.480). Target effects were likewise unsupported when non-optimized and optimized crRNAs were analyzed separately (all asymptotic *p* ≥ 0.107, all PB *p* ≥ 0.243), and optimized-versus-non-optimized contrasts within individual targets were nonsignificant for editing, biallelic editing, and OOF editing (Figure S10). One chr 14 chimerism contrast was significant (*p* = 0.047) but occurred without a supported global optimization × target interaction.

#### 3.3.5 Residual heterogeneity is predominantly line-associated, with generally stable model refits

Variance components were generally negligible at transformation level and small for transformation × target, whereas line-level heterogeneity was pronounced for editing, biallelic editing, and especially chimerism. For OOF editing line-level heterogeneity was lower and for allele bias it was negligible (Table S6). Similar patterns occurred in the *ttLbCas12a* subset. Leave-one-transformation-out refits generally supported the directional robustness of the main construct- and heat-associated effects, whereas sign changes occurred predominantly for target-specific coefficients. All planned refits converged except four of 11 biallelic-editing refits. Construct-associated estimates were less stable for chimerism and allele bias, consistent with the weaker or calibration-dependent inference for these outcomes.

### 3.4 Mutation patterns are dominated by small, cleavage site-centered deletions

Deletions dominated edited alleles (92.9–98.5%). Insertions occurred only in ABA-Cas without heat treatment (3.5%), and mixed InDels remained rare (1.5–7.1%; Table S7). Across all four target sites, deletions ranged from 1–141 bp but were typically small (median 9 bp, IQR 7–10 bp, central 90% 4– 15 bp; Table S8). Median size varied only from 8–9 bp among construct × heat groups, whereas the targets on chr 2/14 tended to show larger deletions than targets on chr 4/9 (Figure 4D; Figure S11). No consistent allelic differences were apparent (Figure S12). Deletion breakpoints clustered around the predicted Cas12a cleavage region, frequently spanning the non-target strand break while retaining the PAM-distal target strand break (Figure S13). Recurrent products varied among conditions without a clear construct- or heat-specific pattern (Figure S14). Most pure-deletion alleles retained ≤ 16 contiguous PAM-proximal spacer nucleotides (93.9%). 4.2% retained 17–18 nt, 1.6% ≥ 19 nt, and 0.2% disrupted the PAM (Figure S15).

#### 3.4.1 Validation of automated mutation characterization

The automated mutation-calling workflow used to derive deletion coordinates for analyses of deletion position relative to the expected cleavage site, recurrent deletion products, and recognition-site retention or loss was evaluated using an independent synthetic hold-out dataset, with additional method agreement assessed against manual experimental annotations. Primary automated calls were obtained for 98.02–98.75% of observations across the development, hold-out, and experimental datasets (Table S9). Multiclass accuracy, reflecting correct discrimination among WT, deletion, insertion, and mixed InDel classes, was 83.04% (Cohen’s κ = 0.773) in the independent synthetic hold-out dataset, while agreement with manual annotations in the experimental dataset was 97.81% (κ = 0.957). Edited-versus-WT accuracy was 98.99% and 99.74%, respectively. Deletion sizes were reproduced exactly for 100% and 98.06% of comparable deletions, while sequence-equivalent breakpoint agreement reached 88.70% and 100%, and recognition-site classifications agreed in 99.13% and 100% of cases, respectively (Tables S10, S11).

## 4 Discussion

This study separates editing efficiency from genotype quality in multiplex *Lb*Cas12a editing of hybrid poplar. While *ttLbCas12a*-based constructs and heat treatment were associated with the strongest increases in biallelic editing efficiency, 3′-extended crRNAs provided a more modest additional benefit, suggesting that guide optimization can further improve editing under conditions supporting high Cas12a activity. In contrast, the recovery of predicted KO-like genotypes depended not only on editing efficiency, but also on repair outcomes, reading-frame consequences, and the ability to obtain biallelic OOF combinations across all target loci.

### 4.1 Balancing heat treatment and selection during primary transformant recovery

Cas12a-mediated genome editing is temperature-sensitive in plants, and elevated-temperature treatments have been shown to increase the activity of several Cas12a variants, including *Lb*Cas12a, in plant systems such as poplar (Malzahn et al. 2019; An et al. 2020). However, the benefit of heat treatment depends on when it is applied during regeneration. Heat exposure should occur early enough to promote editing in the cell lineages that give rise to regenerated shoots, particularly in species with long generation times where primary transformants often constitute the final experimental material rather than an intermediate generation for subsequent segregation (Ding et al. 2020). Conversely, treatment should not be so early or intense that it restricts regeneration, which is already a major bottleneck in many woody species (Wu and Wallner 1983; Castander-Olarieta et al. 2018; Maharjan et al. 2025).

In our workflow, the selected heat-treatment window was compatible with regeneration and antibiotic selection, suggesting that Cas12a-activating temperature exposure can be incorporated without evident impairment of shoot development when applied at an appropriate developmental stage. Consistent with previous reports in other plant systems, the selection regime affected the practical regeneration process, with hygromycin reducing escape frequency but slowing shoot development compared with kanamycin, illustrating a practical trade-off between selection stringency and the rate of shoot regeneration (Rashid 2017; Ontiveros-Cisneros et al. 2022). The higher transgenic recovery observed for the hygromycin-utilizing system further indicates that selection conditions can substantially influence the composition of the regenerated transformant population, although marker system and construct were not independently separated in the present design. The elevated co-cultivation temperature was incorporated to support T-DNA transfer efficiency. However, its independent contribution to transgenic recovery could not be separated from the subsequent heat-treatment component in the present design. Together, these observations support a practical balance between Cas12a-activating heat exposure, regeneration capacity, and effective selection during recovery of primary transformants.

### 4.2 *ttLbCas12a*-based constructs and heat treatment enhance multiplex biallelic editing

Engineered *Lb*Cas12a variants, including tt*Lb*Cas12a carrying the D156R mutation, show improved editing activity at plant-compatible temperatures compared with WT *Lb*Cas12a and have enabled efficient multiplex editing in plants, including stably transformed and regenerated poplar lines (Schindele and Puchta 2020; Zhang et al. 2023). Consistent with these studies, *ttLbCas12a*-based constructs showed high multiplex editing performance in the present study. Importantly, this strategy was applied to a four-locus paralogous target system in heterozygous hybrid poplar, enabling recovery of individual primary transformants carrying biallelic edits across all four *PtaABA8ox4* loci. Such complete editing is particularly relevant for KO-oriented applications, where multiple paralogs and both parental alleles must be modified within the primary generation. Although previous studies reported target-dependent variation in Cas12a-mediated editing (Schindele and Puchta 2020; Zhang et al. 2023), no statistically supported target effect was detected across the four *PtaABA8ox4* loci in the present study. Importantly, the focused analysis of tt*Lb*Cas12a-utilizing approaches likewise provided no consistent evidence that target-specific responses to crRNA optimization masked such differences in the full dataset. Because target identity and crRNA sequence were intrinsically linked, however, the contribution of the target sequence itself cannot be separated from the corresponding crRNA context. Beyond target-level differences, allelic sequence variation around the protospacer could likewise affect Cas12a activity and repair outcomes through local sequence context (Lageix et al. 2024; Nalefski et al. 2024). However, monoallelic editing showed no clear systematic preference for the *P. tremula*-derived A1 or *P. alba*-derived A2 allele, and descriptive A1/A2 deletion patterns likewise revealed no pronounced allele-specific repair differences. Thus, the allelic variation present at the four *PtaABA8ox4* targets did not produce an obvious overall asymmetry in editing or repair outcomes under the conditions tested.

Heat treatment provided an additional improvement in editing performance, with particularly clear gains in biallelic and full-biallelic editing. These findings indicate that elevated temperature remained beneficial even in a workflow based on the temperature-tolerant tt*Lb*Cas12a variant. This is consistent with previous observations in *A. thaliana*, where tt*Lb*Cas12a showed higher activity at 28 °C than at 22 °C (Schindele and Puchta 2020). The broadly positive heat response therefore supports heat treatment as an effective additional workflow component. (An et al. 2020) reported only a modest heat-associated increase for *Lb*Cas12a in poplar. However, direct quantitative comparison is limited by differences in temperature regime, including the additional elevation of co-culture temperature and the higher treatment temperature used here (37 °C vs. 28 °C), which may have contributed to the stronger heat-associated response, as well as by differences in regeneration context, and target architecture. Because RNA secondary structures can respond to temperature in plants, temperature-dependent crRNA folding could also contribute to variation in editing performance across temperature regimes (Su et al. 2018; Creutzburg et al. 2020). Although RNAfold predicted some temperature-dependent changes, the major structural features of the crRNAs were largely retained between 22 °C and 37 °C. In particular, crRNA1 maintained pronounced spacer-associated self-pairing despite increased editing under heat treatment, indicating that altered crRNA folding alone cannot explain the heat-associated gain. Thus, the heat-associated response is consistent with enhanced Cas12a activity, while potential contributions of crRNA folding cannot be excluded (Malzahn et al. 2019; An et al. 2020).

The ABA-Cas versus ABA-ttCas comparison was confounded with selection marker. Selection conditions can influence T-DNA copy-number distribution and the expression of co-integrated transgenes (Zhou et al. 2013). However, ABA-Cas transformants, which showed lower editing efficiency, were recovered under the more stringent HPTII/hygromycin regime, which could be expected to favor events with sufficient T-DNA dosage and transgene expression. Thus, although the selection regime may have influenced the magnitude of the observed differences, it does not readily explain their overall direction, with *ttLbCas12a*-based transformants consistently outperforming *LbCas12a*-based transformants. Nevertheless, nuclease and selection-marker effects cannot be fully separated. Because transgene copy number and expression data were not determined, potential contributions of T-DNA dosage, insertion context, or Cas12a/crRNA expression to the observed line-to-line variation remain unresolved. Fully crossed construct designs will therefore be required to isolate the contribution of the nuclease variant, while additional transgene characterization would be needed to resolve mechanistic sources of line-to-line variation. Together, these results support the combination of *ttLbCas12a*-based constructs and heat treatment as an effective strategy for multiplex biallelic editing in hybrid poplar, here enabling regeneration of primary transformants edited across all four paralogous *PtaABA8ox4* target loci.

### 4.3 crRNA 3’-tailing is compatible with efficient editing but its independent benefit remains unresolved

Cas12a editing performance depends on crRNA and target context, while 3′ extensions may improve activity by altering crRNA secondary structure (An et al. 2020; Creutzburg et al. 2020; Li et al. 2023a; Ren et al. 2023; Zhang et al. 2023; Nguyen et al. 2024). Accordingly, our optimized crRNAs carried short 3′ tails designed to improve predicted crRNA architecture. RNAfold predictions indicated reduced spacer-associated self-pairing in crRNA1.opt, potentially increasing spacer accessibility, whereas crRNA2.opt showed increased base-pairing probability in the pseudoknot-forming region, which may stabilize the Cas12a-compatible crRNA architecture and reduce susceptibility to competing alternative folds involving the spacer (Creutzburg et al. 2020). 3′-tailed crRNAs showed a consistent numerical tendency toward improved editing, but neither the full-dataset comparisons nor the focused tt*Lb*Cas12a analysis statistically resolved an independent benefit of 3′-tailing.The already high editing efficiency of non-tailed crRNAs, particularly under heat treatment, may have limited the scope for further improvement, while the absence of an ABA-Cas.opt construct prevented testing whether tailing is more beneficial in a lower-activity *Lb*Cas12a background. Interpretation was further limited by the small number of target-paired crRNA sets, as the benefit of 3′-tailing may depend on guide context and the underlying level of Cas12a activity. Guide context may also include tail–target complementarity, because complementary 3′ extensions can alter Cas12a target recognition and cleavage kinetics (Strohkendl et al. 2018; Naqvi et al. 2022). Here, limited allele-specific tail–target complementarity did not translate into clear allelic bias. Thought, subtler effects on guide activity or cleavage dynamics cannot be excluded. Moreover, the divergent RNAfold-predicted structures were not accompanied by correspondingly distinct editing performance, consistent with the limitations of secondary-structure prediction for estimating Cas12a crRNA functionality reported by Creutzburg et al. (2020). Nevertheless, the high performance of ABA-ttCas.opt demonstrates that 3′-tailed crRNAs are compatible with efficient multiplex editing in stably transformed and regenerated poplar lines. Because short tails can be encoded directly in the crRNA expression cassette, 3′-tailing provides a simple guide-engineering option that may be useful when spacer choice is constrained or as an additional optimization layer. Future studies should test tailed and non-tailed crRNAs across larger target-paired designs with matched construct contexts and directly assess crRNA processing, abundance, and activity.

### 4.4 Increased editing is not consistently accompanied by increased chimeric biallelic outcomes

Somatic genotypic heterogeneity, here operationally termed chimerism, is a major challenges in genome editing of woody perennials, where primary transformants are often used directly and chimeric genotypes cannot readily be resolved by segregation (Ding et al. 2020; Cui et al. 2021; Nanasato et al. 2021). Chimerism has been reported for both Cas9- and Cas12a-based editing in poplar and can arise from multicellular shoot origin or from delayed nuclease activity during regeneration (Ding et al. 2020; Li et al. 2023a; Zhang et al. 2023). Increased Cas12a activity could either reduce chimerism through earlier editing or promote additional somatic variants through persistent cleavage during regeneration, particularly because Cas12a cleavage can retain PAM-proximal recognition features that permit re-cleavage (Zetsche et al. 2015; Gong et al. 2021). High editing efficiency can likewise coexist with substantial chimerism in poplar, as reported by Li et al. (2023a) using a different Cas12a variant. In the present study, chimerism among biallelically edited targets did not consistently increase with editing efficiency across constructs and heat treatments. We hypothesize that initial editing frequently reduced subsequent target recognition: most pure-deletion alleles retained no more than 16 contiguous PAM-proximal spacer nucleotides, likely reducing susceptibility to efficient re-cleavage (Strohkendl et al. 2018; Naqvi et al. 2022; Ren et al. 2023). This may limit continued somatic mutagenesis and favor stability of already edited alleles during clonal propagation. Chimerism nevertheless showed descriptive construct-specific variation, which may reflect differences in the timing and persistence of editing activity. Guide– target interactions and complementary 3′ crRNA extensions can alter Cas12a target recognition and cleavage kinetics (Strohkendl et al. 2018; Naqvi et al. 2022), potentially shifting editing to later stages of regeneration. Guide-dependent mutation outcomes may also affect how rapidly edited alleles lose susceptibility to re-cleavage, as crRNA guide length can alter Cas12a-induced deletion-size distributions even at the same target site (Negishi et al. 2020). Together with multicellular shoot origin, these processes could contribute to residual chimerism. However, endpoint genotyping cannot distinguish among these mechanisms, and Sanger sequencing may additionally miss low-frequency chimeric sectors. Deeper sequencing would therefore be required for more sensitive quantification, while secondary regeneration may provide a practical strategy to resolve valuable chimeric lines, as demonstrated for CRISPR/Cas9-edited poplar (Ding et al. 2020).

### 4.5 Mutation patterns and reading-frame consequences

Because Cas12a-induced mutation spectra can depend on nuclease ortholog, target/crRNA context, and local sequence environment, we examined whether the optimization strategies also altered deletion patterns (Malzahn et al. 2019; Lageix et al. 2024; Li et al. 2023a). Mutations were dominated by small deletions, consistent with previous Cas12a studies in poplar and other woody species (Jia et al. 2022; Ren et al. 2023; Das et al. 2025; Zhang et al. 2023; Zahn et al. 2026). Given the predominance of deletions and the high accuracy of deletion-size and breakpoint assignment, the automated mutation-characterization workflow was well suited for analyzing Cas12a-induced deletion patterns. The approach should be transferable to comparable Sanger sequencing datasets that are not readily compatible with standard amplicon-based analysis tools, provided that target references and cleavage sites are known. Deletions frequently span the non-target strand cleavage position while retaining the PAM-distal target strand cleavage position, compatible with processing of Cas12a-generated staggered ends during end joining (Lageix et al. 2024). No consistent heat- or construct-associated shift in deletion size or positional patterns was apparent, suggesting that the conditions increasing editing efficiency did not substantially redirect the dominant mutation profile. The absence of a clear temperature-associated shift is consistent with Lageix et al. (2024), whereas nuclease-dependent effects on Cas12a mutation profiles appear context-dependent, with both distinct and broadly similar deletion patterns reported across nuclease comparisons (Malzahn et al. 2019; Li et al. 2023a). Accordingly, the similar profiles of WT *Lb*Cas12a and tt*Lb*Cas12a provide no indication that the D156R modification substantially altered the dominant deletion pattern under the conditions tested. Descriptively, deletion patterns varied among target/crRNA contexts and were more similar between targets sharing the same crRNA, consistent with target- and crRNA-associated mutation patterns reported in perennial species (Ren et al. 2023; Zhang et al. 2023; Das et al. 2025; Zahn et al. 2026). Because target and crRNA identity were linked in the present design, however, their individual contributions cannot be separated, and the limited number of recovered pure-deletion alleles, particularly in low-efficiency treatment groups, warrant cautious interpretation.

The deletion-dominated mutation spectrum also illustrates why high editing efficiency does not necessarily translate into KO-like genotype recovery. Cas12a-induced deletions varied in length and included both frame-disrupting and frame-preserving outcomes. Accordingly, increases in biallelic editing were not mirrored proportionally by biallelic OOF recovery: although multiple lines carried biallelic OOF edits at several loci, none combined such outcomes across all four *PtaABA8ox4* targets. Given the limited number of recovered lines, the compounded requirement for biallelic OOF outcomes across all four loci, and the absence of a clear target-specific bias toward OOF or in-frame outcomes, this does not provide evidence for negative selection against complete *PtaABA8ox4* frameshift genotypes. Total editing and biallelic editing rates alone may therefore overestimate KO-like recovery when the functional consequences of individual mutations are not considered. For knockout-oriented applications, target selection should prioritize functionally critical sequence regions, while paired crRNAs may provide an alternative strategy for generating larger disruptive deletions spanning essential motifs or domains (An et al. 2020). Because OOF classification was based solely on predicted reading-frame consequences, these genotypes should be regarded as predicted KO-like rather than experimentally confirmed loss-of-function lines.

## 5 Conclusions

In conclusion, *ttLbCas12a*-based constructs combined with heat treatment during co-culture and regeneration substantially improved multiplex biallelic genome editing in heterozygous hybrid poplar and enabled recovery of primary transformants carrying biallelic edits across all four targeted *PtaABA8ox4* loci. crRNA 3′-tailing was compatible with highly efficient editing and may provide an additional low-cost design lever, but its independent contribution remains to be validated in larger, target-paired datasets. However, the recovery of biallelic edits alone was insufficient to define KO-like genotypes, because genotype usability also depended on somatic uniformity, mutation profiles, reading frame consequences, and target position. This workflow provides a practical basis for multiplex Cas12a-mediated editing in poplar and highlights evaluation criteria that are particularly relevant for highly heterozygous and duplicated woody genomes.

## CRediT authorship contribution statement

**Virginia Zahn:** Conceptualization, Methodology, Software, Validation, Formal analysis, Investigation, Data curation, Visualization, Writing – original draft, Writing – review & editing. **Alexander Fendel:** Conceptualization, Methodology, Investigation, Writing – review & editing. **Matthias Fladung:** Conceptualization, Writing – review & editing, Funding acquisition. **Tobias Bruegmann:** Conceptualization, Methodology, Formal analysis, Resources, Supervision, Project administration, Funding acquisition, Writing – review & editing.

## Acknowledgements

We are grateful for the excellent laboratory assistance of Susanne Jelkmann, Katrin Groppe, Cid Redlin, and Ian Volkmann.

## Data Availability

All data generated and analyzed in this study are provided in Supplementary Data S1–S6. Data S1 contains vector maps in GenBank (.gb) format. Data S2 contains detailed outputs, sequence files, and raw data used for automated genotype calling and validation. Data S3 contains the experimental raw data. Data S4 contains the plot data underlying the figures. Data S5 contains detailed outputs from the GLMM analyses. Data S6 contains the detailed mutation-pattern analyses. The R scripts used for statistical analyses, automated mutation characterization, validation, and data visualization are available via Zenodo (Zahn 2026). The plant material generated in this study will be made available to other researchers upon reasonable request, subject to availability at the time of request.

